# FPD: A comprehensive phosphorylation database in fungi

**DOI:** 10.1101/058867

**Authors:** Youhuang Bai, Bin Chen, Yincong Zhou, Silin Ren, Qin Xu, Ming Chen, Shihua Wang

**Author notes:** These authors contributed equally to this work (Youhuang Bai, Bin Chen). To whom correspondence should be addressed: Prof. Ming Chen, or Prof. Shihua Wang.

## Abstract

Protein phosphorylation, one of the most classic post-translational modification, plays a critical role in the diverse cellular processes including cell cycle, growth and signal transduction pathways. However, the available information of phosphorylation in fungi is limited. Here we provided a Fungi Phosphorylation Database (FPD) that comprises high-confidence in vivo phosphosites identified by MS-based proteomics in various fungal species. This comprehensive phosphorylation database contains 62,272 non-redundant phosphorylation sites in 11,222 proteins across eight organisms, including *Aspergillus flavus, Aspergillus nidulans, Fusarium graminearum, Magnaporthe oryzae, Neurospora crassa, Saccharomyces cerevisiae, Schizosaccharomyces pombe* and *Cryptococcus neoformans*. A fungi-specific phosphothreonine motif and several conserved phosphorylation motif were discovered by comparatively analyzing the pattern of phosphorylation sites in fungi, plants and animals.

Database URL: http://bis.zju.edu.cn/FPD/index.php

## Introduction

Posttranslational modification (PTM) in particular plays a significant role in a wide range of cellular processes, and provides an additional layer to determine the kinetics and cellular plasticity through the dynamical cell signaling networks. Phosphorylation, mainly occurs on three types of amino acids (serine, threonine and tyrosine), is a very common mechanism for regulating the activity of enzymes to monitor events and initiate appropriate responses. Recently, Greig *et al.* show that changes of phosphorylation status of the fork-head family transcription factor Fkh2 drives the pathogenic switch in the opportunistic human fungal pathogen *Candida albicans* (1).

As the model organism, yeast attracted special attentions from the fungi community and it takes continuous effort to improve the quality of connection between the dynamic phosphorylation event and function of proteins (2,3). Although the phosphoproteomics techniques in yeast developed quickly, phosphoproteome analysis in other fungi is still limited. The Xiong Y *et al.* identified and quantified 1, 942 proteins with 5,882 phosphorylation sites with high confidence from two replicates in *Neurospora crassa* (4). Selvan *et al.* identified 1089 phosphopeptides derived from 648 proteins including about 45 kinases in *Cryptococcus neoformans* (5). Recently we analyzed the *Aspergillus flavus* phosphoproteome and identified 598 high-confidence phosphorylation sites in 278 phosphoproteins (Ren S. *et al.,* unpublished). And a great deal of effort has been made to identify phosphorylation events in other fungi, such as *Aspergillus nidulans* (6), *Alternaria brassicicola* (7), and *Botrytis cinerea* (7,8). With the accumulation of phosphoproteome studies in fungi, a large number of phosphorylation events and sites were identified, while data maintenance and sharing became increasingly challenging. These numerous phosphorylation data in fungi may contribute to expand the understanding of molecular mechanisms and functional roles for phosphorylation in fungal communities.

Until now, several online databases collected and integrated numerous phosphorylated substrates with their sites from different species, such as PhosphositePlus (9), Phospho.ELM (10), PHOSIDA (11), PhosphoPep (12) and P3DB (13). Recently Xue Y. lab built three databases of protein phosphorylation sites in prokaryotes (14), plants (15), animals and fungi (16). However, the dbFAF database had only collected the phosphorylation sites in two fungi *S. cerevisiae* and *S. pombe.* Besides, several databases mainly focused on phosphorylation in specific species, for example, PhosphoGRID (*S. cerevisiae*) (17), PhosPhAt (*Arabidopsis thaliana*) (18), and HPRD (Human) (19). To our knowledge, only a limited proportion of the identified phosphoproteins and phosphosites in fungi were covered by public database.

In this study, we provide a comprehensive collection of 62,272 non-redundant phosphorylation sites from 11,222 proteins that were collected from literatures and database. The phosphoproteins were from eight fungal organisms, including *Aspergillus flavus, Aspergillus nidulans, Fusarium graminearum, Magnaporthe oryzae, Neurospora crassa, Saccharomyces cerevisiae, Schizosaccharomyces pombe* and *Cryptococcus neoformans.* These data were collected in Fungal Phosphorylation Database (FPD, https://bis.zju.edu.cn/FPD/index.php). By examining the motif of phosphorylation events in three kingdoms, the conservation of phosphorylation motifs was discovered. Taken together, the FPD database could serve as a comprehensive protein phosphorylation data resource for further studies in fungi.

## Methods

The FPD is a relational database built on MySQL server. The web application runs on an Apache version 2.4.7 server; in-house developed PHP scripts provide data retrieval. The web interface is based on PHP and CSS scripts. The FPD is publicly available at http://bis.zju.edu.cn/FPD/index.php.

We searched the PubMed with multiple keywords: (“phosphoproteomics” OR “phosphoproteomic” OR “phosphoproteome”) and (“fungal” OR “fungi”). All retrieved articles were carefully curated. Fungal protein sequences were collected from Uniprot database. The identified phosphorylated proteins, peptides and sites from the supplementary materials published together with these manuscripts if available (Supplementary Table S1). For each species, we mapped corresponding phosphorylated peptides to the protein sequences, and the phosphorylation sites were exactly pinpointed by using in-house Perl scripts. Besides curation from literatures, phosphorylation sites of *S. cerevisiae* and *S. pombe* in public databases dbPAF were integrated into our database. The detailed annotations of phosphoproteins were retrieved from Uniprot.

To identify the conserved phosphosite motif, we analyzed the motifs of all phosphorylated sites in plants, animals and fungi. All phosphorylated sites in four reprehensive plants (*Arabidopsis thaliana, Medicago truncatula, Oryza sativa* and *Zea mays*) were downloaded from dbPPT. These data in five animals (*Rattus norvegicus, Drosophila melanogaster, Caenorhabditis elegans, Homo sapiens* and *Mus musculus*) were downloaded from dbPAF. Phosphorylated peptides in length of 13 with central characters of S/T/Y residues were prepared as the foreground data set, while non-phosphorylated peptides in the same proteins were regarded as the background data set. The phosphorylation motifs were calculated for three types of residues, respectively. By submitting phosphorylation peptides and non-phosphorylation peptides to Motif-X (http://motif-x.med.harvard.edu/motif-x.html), all significant motifs for each species were gathered. The data of *Homo sapiens* and *Mus musculus* were excluded in phosphoserine motif analysis because the data size is too large to get result under the default parameter setting of Motif-X.

## Database usage

To provide convenient services for sharing the phosphorylation information to the fungi community, FPD database was designed to contain browse and search functions with various options in a user-friendly manner. The browse function is developed for organism-based querying among the eight fungi species. The species are presented in an evolutionary tree on the ‘BROWSE’ page, where user could click the image or organism name for one species to browse all phosphoproteins. Using the important filamentous fungi *Aspergillus flavus* as an example, the workflow for browsing is shown in Fig 1. All the identified phosphoproteins of *A. flavus* are listed as a table containing four type of basic information, including unique FPD_ID, Uniprot id, gene name and organism. User could click the FPD_ID entry for the protein of interest to view the details of the phosphoprotein. This page contains more information, including database links, sequence and the position of phosphorylation sites in proteins. The phosphorylation sites are presented with position number, peptide with 7 amino acids upstream and downstream of the modified residue.

**Figure 1.**
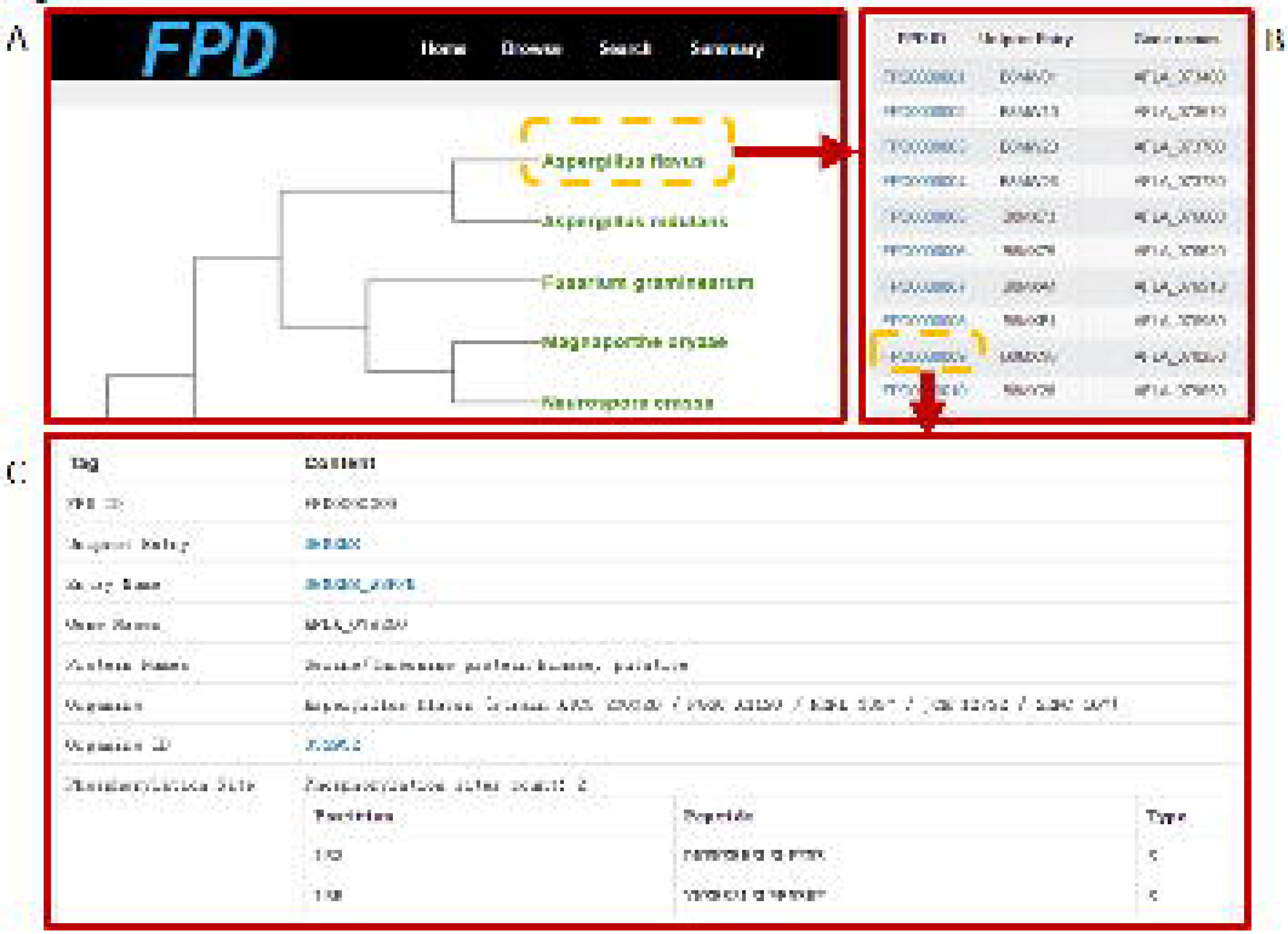
Snapshots of browser function in the FPD database. (A) The tree of eight fungi species which contain identified phosphorylation sites. (B) The list of identified phosphoprotein in *Aspergillus flavus*. (C) The detailed information of AFLA_078250 protein, including the phosphorylation information.

To provide convenient querying services, different search options are implemented in “Search” page of FPD database. A search option is provided, which allows user to perform keyword-based search in multiple specific areas. Take AFLA_046360 in example, Fig. 2A is shown by searching the keyword “*Aspergillus flavus*” in the “Organism” area, “AFLA_046360” in the “gene name” area, and “carboxylase” in the “Protein Names”. After submitting the query, the results will be shown in a tabular format with basic information, and then detail information is available by clicking the FPD_ID entry. In addition, a batch search mode, as shown in Fig. 2B, is also provided for retrieving multiple phosphoproteins with a list of keywords.

**Figure 2.**
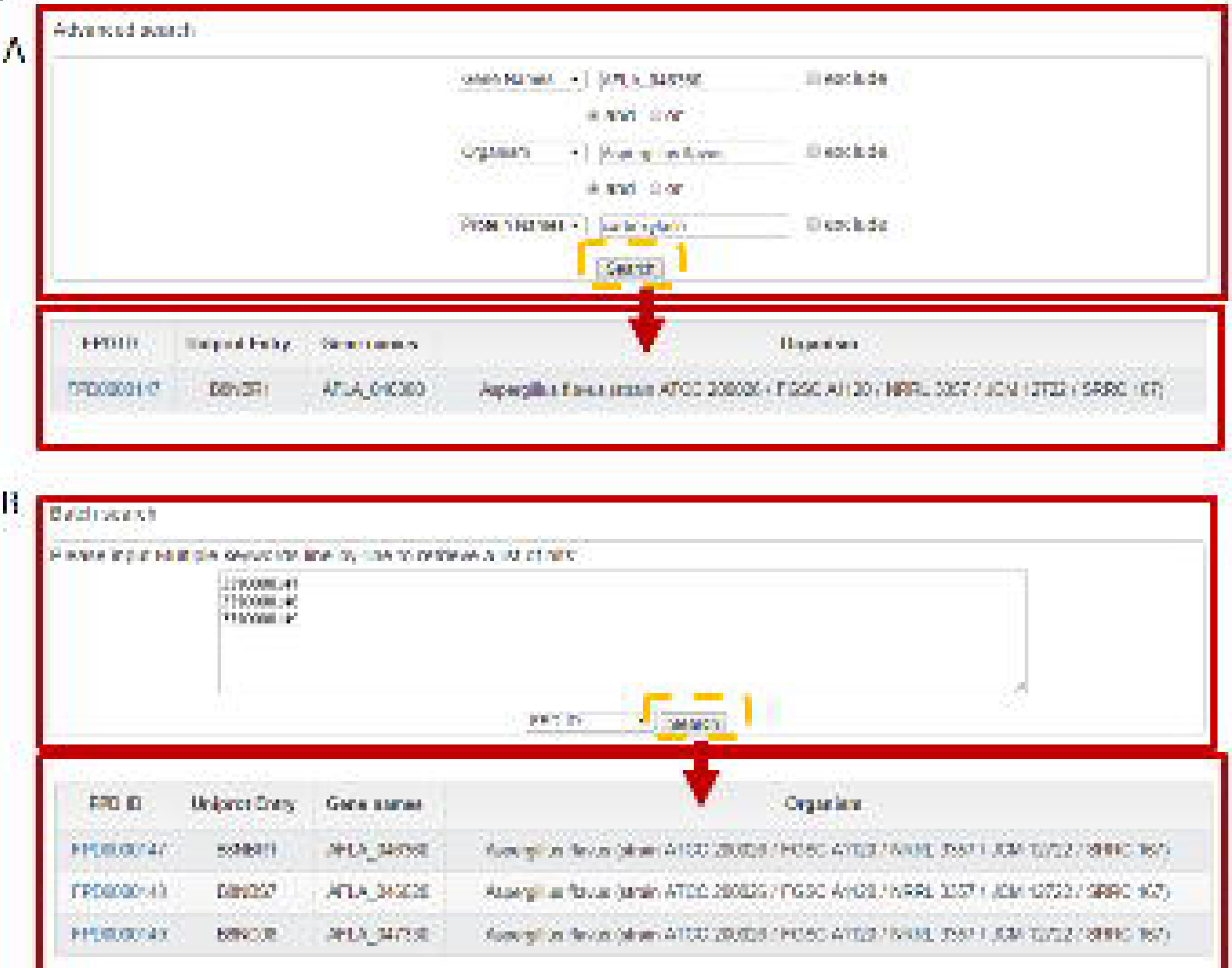
Snapshots of search function in the FPD database. (A) FPD can be queried with one or multiple keyword(s) with multiple operators, such as “AFLA_046360” in the “gene name” area. (B) Batch search allows users to search multiple keywords by one click.

## Conclusion and future direction

As one of the most important PTMs, protein phosphorylation was involved in almost all aspects of biology processes. While the collection of these information in fungi is less, except for yeast. Phosphoproteome in fungi species was extensively studied in recent years, and tens of thousands of phosphorylation sites of diverse fungi were identified. So a comprehensive database of fungi phosphorylation information is indeed in demand.

In this study, FPD database provides 62,272 identified phosphorylation sites in 11, 222 phosphoproteins in eight fungi ranged from Agaricomycotina, Taphrinomycotina, Saccharomycotina to a large group of Pezizomycotina (Table 1). The proportions of phosphoserine (pS), phosphothreonine (pT) and phosphotyrosine (pY) were 73.20%, 22.98% and 3.82%, respectively. The number of phosphosites per phosphoprotein is as high as 9.8 in *Saccharomyces cerevisiae,* while this value is range from 2.15 to 3.73 in the other seven fungi. It is the fact that phosphorylation events discovered in *Saccharomyces cerevisiae* are more abundant than that in rest seven fungi. Furthermore, in the foreseeable future, more and more experimental and computational studies of phosphoproteomic data will lead to the boom of phosphorylation sites in fungi.

In order to find that whether the sequence motifs for phosphosites exists or not in all three kingdoms, a comparative analysis of all phosphorylation pattern was done by using motif-X algorithm. The phosphorylation events in plants and animals were also collected. These data were integrated with fungi phosphosites in FPD. By submitting phosphorylation peptides and non-phosphorylation peptides to Motif-X, all significant motifs were retrieved and analyzed (Fig. 3A, Fig. 4A, and Supplementary Table S4). For serine phosphorylation, three motifs emerged in phosphoserine sites in 15 species analyzed: “pS-D-x-E”, “pS-P” and “R-x-x-pS” (where x represents any amino acid) (Fig. 3B–D, Supplementary Table S2). In HPRD database (http://www.hprd.org/), “pS-D-x-E” motif is classified as Casein kinase (CK) II substrate motif, which might be substrates of CKII. In plants and animals, it was proposed that “pS-P” motif might be associated with activity of MAPKs and cyclin dependent kinase (CDK), while “R-X-X-pS” is PKA kinase substrate motif, which might be recognized by CPKs and CaMKs. In addition, several motifs were found to be absent in one specific kingdom while present in other kingdoms. For example, “pS-P-x-R” motif is recognized as CDK1, 2, 4, 6 kinase substrate motif, which is absent from the all fungi species analyzed but present in both plants and animals.

**Figure 3.**
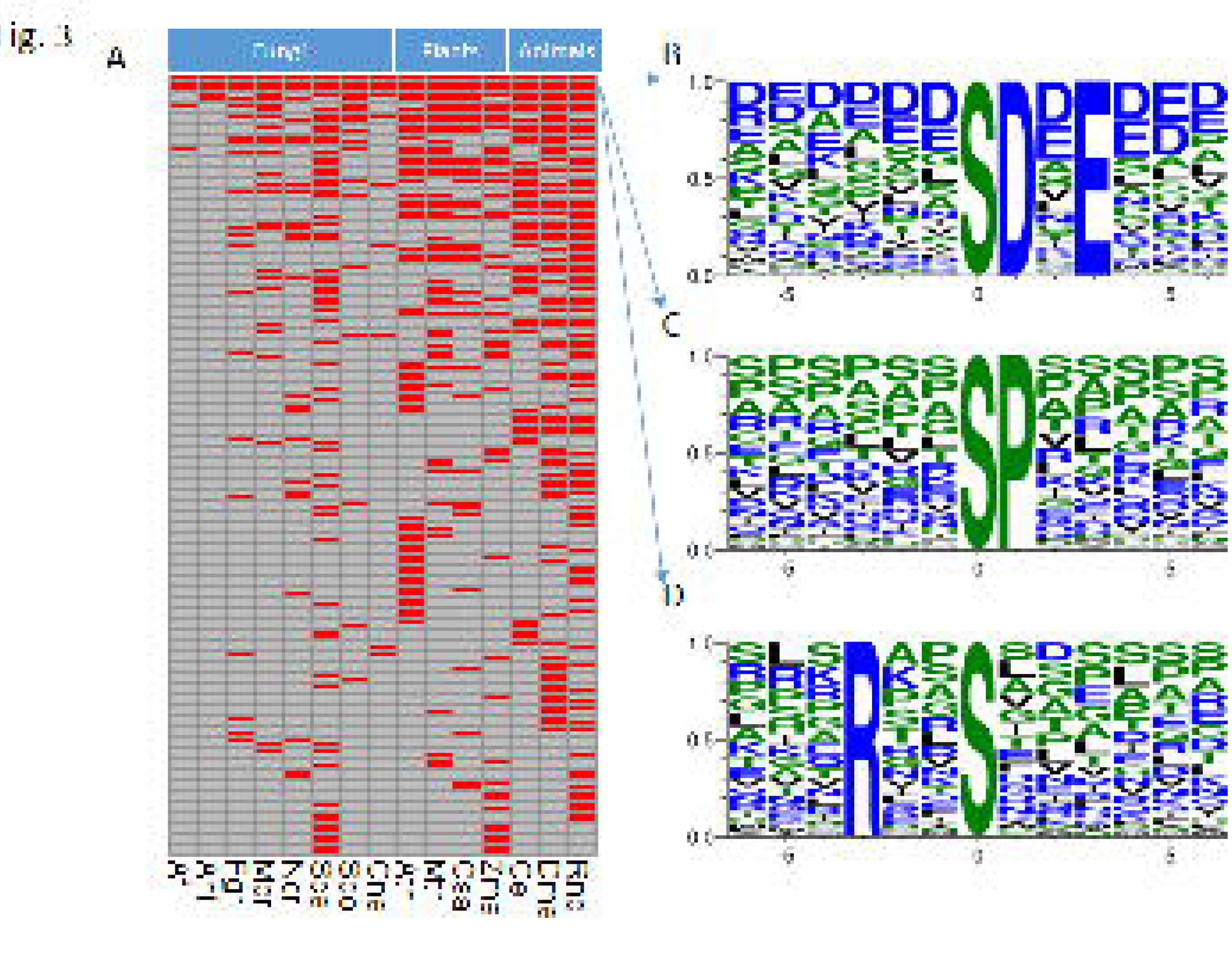
Comparative analysis of phosphoserine motifs in fungi, plants and animals. (A) The heatmap of each significant pS motif in the species analyzed (Afl, *Aspergillus flavus*; Ani, *Aspergillus nidulans*; Fgr, *Fusarium graminearum*; Mor, *Magnaporthe oryzae*; Ncr, *Neurospora crassa*; Sce, *Saccharomyces cerevisiae*; Spo, *Schizosaccharomyces pombe*; Cne *Cryptococcus neoformans*; Rno, *Rattus norvegicus*; Dme, *Drosophila melanogaster*; Cel, *Caenorhabditis elegans*; Ath, *Arabidopsis thaliana*; Mtr, *Medicago truncatula*; Osa, *Oryza sativa*; Zma, *Zea mays*) (B) weblogo of “pS-P” motif. The position 0 represents the modified residues. (C) weblogo of “pS-D-x-E” motif. (D) weblogo of “R-x-x-pS” motif.

**Figure 4.**
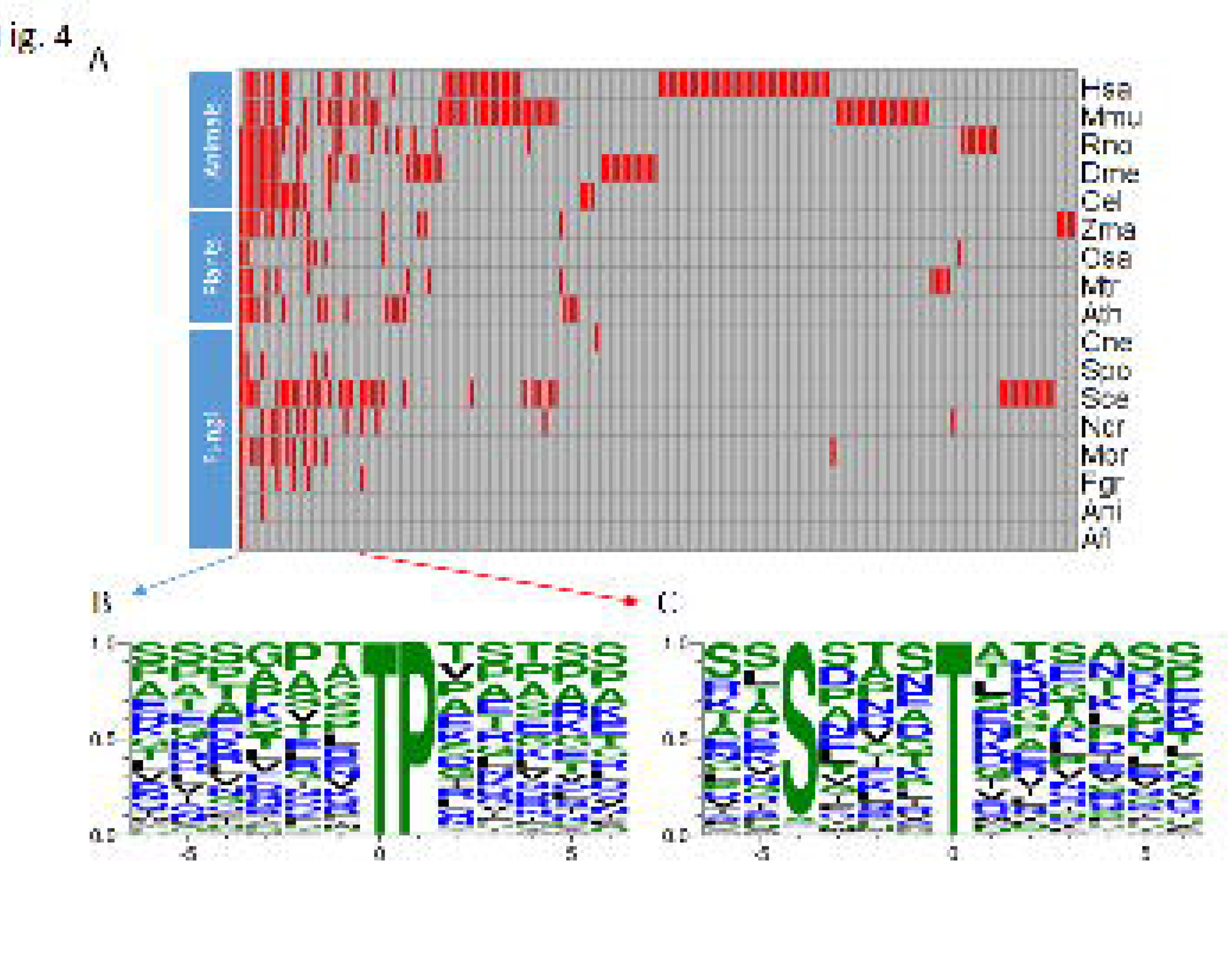
Comparative analysis of phosphothreonine motifs in fungi, plants and animals. (A) The heatmap of each significant pT motif in the species analyzed (Afl, *Aspergillus flavus*; Ani, *Aspergillus nidulans*; Fgr, *Fusarium graminearum*; Mor, *Magnaporthe oryzae*; Ncr, *Neurospora crassa*; Sce, *Saccharomyces cerevisiae*; Spo, *Schizosaccharomyces pombe*; Cne *Cryptococcus neoformans*; Rno, *Rattus norvegicus*; Dme, *Drosophila melanogaster*; Cel, *Caenorhabditis elegans*; Mmu, *Mus musculus*; Has, *Homo sapiens*; Ath, *Arabidopsis thaliana*; Mtr, *Medicago truncatula*; Osa, *Oryza sativa*; Zma, *Zea mays*). (B) weblogo of “pT-P” motif. The position 0 represents the modified residues. (C) weblogo of fungi-specific motif “S-x-x-x-pT”.

For pT site, the “pT-P” motif was found in all most species but not in human and mouse, which is also recognized by MAPKs and CDKs (Fig. 4B, Supplementary Table S3). It’s worth noting that a fungi-specific motif “S-x-x-x-pT” was identified in *F. graminearum, N. crassa, S. cerevisiae* (Fig. 4C). In mammals, the similar motif “S/T-x-x-x-pS/pT” is the consensus pattern of phosphorylation site in GSK3 that is Ser/Thr protein kinase and a key regulator of glycogen metabolism. In previous study, GSK3 orthologs was reported from *F. graminearum* (FGSG_07329) and *S. cerevisiae* (MCK1, RIM11, MRK1, and YGK3) (20,21). It is suggested that the motif “S-x-x-x-pT” might be relevant to activity of GSK3 orthologs in fungi, which needed to be experimental validated. Although most motifs were conserved between fungi, plants and mammals, this data indicated that the fungi-specific motif may play important roles in growth and pathogenicity in fungi.

It is expected that more efforts will be contributed to phosphoproteome studies in fungi and the FPD database will be routinely updated to add more information of phosphorylation in fungi species. Taken together, the FPD database provides a comprehensive protein phosphorylation resource for better understanding of phosphorylation in fungi.

## Supplementary Data

Supplementary data are available at Database Online.

Supplementary Table S1. The data source of the phosphoproteins and phosphosites hosted in FPD.

Supplementary Table S2. The details of phosphoserine motifs in fungi, plants and animals. “1” represents that each motif exists in the species, while “0” represents that motif does not exist in the species.

Supplementary Table S3. The details of phosphothreonine motifs in fungi, plants and animals. “1” represents that each motif exists in the species, while “0” represents that motif does not exist in the species.

Supplementary Table S4. The details of phosphotyrosine motifs in fungi, plants and animals. “1” represents that each motif exists in the species, while “0” represents that motif does not exist in the species.

## Acknowledgements

We thank the authors of the original studies of phosphoproteome in each species in our database. Also thanks to Dr. Yu Xue (Huazhong University of Science and Technology) for providing the phosphorylation datasets in dbPSP, dbPPT and dbPAF, respectively.

## References

1. Greig, J. A., Sudbery, I.M., Richardson, J.P., et al. (2015) Cell cycle-independent phospho-regulation of Fkh2 during hyphal growth regulates Candida albicans pathogenesis. PLoS Pathog, 11, e1004630.

2. Kanshin, E., Kubiniok, P., Thattikota, Y., et al. (2015) Phosphoproteome dynamics of Saccharomyces cerevisiae under heat shock and cold stress. Mol. Syst. Biol., 11, 813.

3. Amoutzias, G. D., He, Y., Lilley, K.S., et al. (2012) Evaluation and properties of the budding yeast phosphoproteome. Mol. Cell. Proteomics, 11, M111 009555.

4. Xiong, Y., Coradetti, S.T., Li, X., et al. (2014) The proteome and phosphoproteome of Neurospora crassa in response to cellulose, sucrose and carbon starvation. Fungal Genet. Biol., 72, 21–33.

5. Selvan, L. D., Renuse, S., Kaviyil, J. E., et al. (2014) Phosphoproteome of Cryptococcus neoformans. Journal of proteomics, 97, 287–295.

6. Ramsubramaniam, N., Harris, S. D., Marten, M. R. (2014) The phosphoproteome of Aspergillus nidulans reveals functional association with cellular processes involved in morphology and secretion. Proteomics, 14, 2454–2459.

7. Davanture, M., Dumur, J., Bataille-Simoneau, N., et al. (2014) Phosphoproteome profiles of the phytopathogenic fungi Alternaria brassicicola and Botrytis cinerea during exponential growth in axenic cultures. Proteomics, 14, 1639–1645.

8. Lineiro, E., Chiva, C., Cantoral, J. M., et al. (2016) Phosphoproteome analysis of B. cinerea in response to different plant-based elicitors. Journal of proteomics. doi, 10.1016/j.jprot.2016.03.019

9. Hornbeck, P. V., Zhang, B., Murray, B., et al. (2015) PhosphoSitePlus, 2014: mutations, PTMs and recalibrations. Nucleic Acids Res., 43, D512–520.

10. Dinkel, H., Chica, C., Via, A., et al. (2011) Phospho.ELM: a database of phosphorylation sites--update 2011. Nucleic Acids Res., 39, D261–267.

11. Gnad, F., Gunawardena, J., Mann, M. (2011) PHOSIDA 2011: the posttranslational modification database. Nucleic Acids Res., 39, D253–260.

12. Bodenmiller, B., Campbell, D., Gerrits, B., et al. (2008) PhosphoPep--a database of protein phosphorylation sites in model organisms. Nat. Biotechnol., 26, 1339–1340.

13. Yao, Q., Ge, H., Wu, S., et al. (2014) P(3)DB 3.0: From plant phosphorylation sites to protein networks. Nucleic Acids Res., 42, D1206–1213.

14. Pan, Z., Wang, B., Zhang, Y., et al. (2015) dbPSP: a curated database for protein phosphorylation sites in prokaryotes. Database, 2015, bav031.

15. Cheng, H., Deng, W., Wang, Y., et al. (2014) dbPPT: a comprehensive database of protein phosphorylation in plants. Database, 2014, bau121.

16. Ullah, S., Lin, S., Xu, Y., et al. (2016) dbPAF: an integrative database of protein phosphorylation in animals and fungi. Scientific reports, 6, 23534.

17. Sadowski, I., Breitkreutz, B. J., Stark, C., et al. (2013) The PhosphoGRID Saccharomyces cerevisiae protein phosphorylation site database: version 2.0 update. Database, 2013, bat026.

18. Zulawski, M., Braginets, R., Schulze, W. X. (2013) PhosPhAt goes kinases--searchable protein kinase target information in the plant phosphorylation site database PhosPhAt. Nucleic Acids Res., 41, D1176–1184.

19. Goel, R., Harsha, H. C., Pandey, A., et al. (2012) Human Protein Reference Database and Human Proteinpedia as resources for phosphoproteome analysis. Molecular bioSystems, 8, 453–463.

20. Andoh, T., Hirata, Y., Kikuchi, A. (2000) Yeast glycogen synthase kinase 3 is involved in protein degradation in cooperation with Bul1, Bul2, and Rsp5. Mol. Cell. Biol, 20, 6712–6720.

21. Wang, C., Zhang, S., Hou, R., et al. (2011) Functional analysis of the kinome of the wheat scab fungus Fusarium graminearum. PLoSPathog, 7, e1002460.

